# Structural financial ambiguity in climate-smart agriculture research: A bibliometric analysis of African knowledge systems

**DOI:** 10.64898/2026.03.02.708951

**Authors:** Pierre Marie Chimi, Guillaine Yonga, Albert Tchopwe Menkamla, Benoit Maralossou, Angele Marie Ngon Dikoume, Laskine Mazak Nguihi, Ulrich D’Assise Mvondo Effa, Joseph Martin Bell, William Armand Mala

## Abstract

Climate-smart agriculture (CSA) dominates African agricultural policy discourse, yet fifteen years post-conceptualization, its transformative potential remains unrealized. Bibliometric analysis of 161 Scopus-indexed publications (2014–2025) reveals exponential field growth (31.3% annually) coupled with on technical dimensions and systemic neglect of financial mechanisms. Network analysis (VOSviewer), semantic mapping and citation bibliometrics expose cognitive oligopolisation—wherein 1.8% of authors generate 45% of output—geographical fragmentation into weakly connected regional clusters, and critical underrepresentation of the vulnerable Sahel. Despite 46.6% of publications addressing economic themes, merely 5.6% rigorously integrate financial analysis with adoption variables; terms including ‘investment’, ‘cost–benefit’ and ‘climate finance’ remain absent from major semantic clusters. The concept of ‘structural financial ambiguity’ is introduced to characterize the maintenance of CSA in operational indeterminacy through academic discourse that substitutes description for actionable financial theorization. Paradigmatic transformation conditions are identified through emerging scholarship employing discrete choice experiments and cost–benefit evaluations to construct requisite knowledge foundations. Findings indicate that without comprehensive theorization of microfinance, digital finance, index-based insurance and payments for environmental services, international climate commitments risk implementation failure due to absence of scientifically validated financial instruments rather than technical solutions.

## Introduction

Agriculture in Africa engages approximately 60% of the working population and contributes substantially to national GDP, yet remains highly vulnerable to climate change impacts [1], [2], [3]. Projections indicate potential production declines of 10–20% by 2050 in sub-Saharan Africa, threatening food security for rapidly expanding populations [4], [5]. Climate-smart agriculture (CSA) has emerged as a key strategy for food system transformation, proposing to reconcile productivity enhancement, climate resilience, and greenhouse gas emissions reduction [6], [7].

The translation of these aspirations into outcomes encounters a fundamental barrier: financing. Despite global commitments made at COP21 in Paris ($100 billion annually for adaptation) and reiterated at subsequent COPs, actual monetary flows toward climate-resilient agriculture in Africa remain inadequate and misallocated [8], [9]. The African continent requires an estimated additional $257 billion in investments by 2030 to achieve sustainable development objectives, with agriculture constituting a significant proportion [10], [11]. This disparity between commitments and outcomes cannot be reduced to technical or resource limitations. Drawing on transition research [12], [13] and institutional economics [14], [15], CSA implementation is framed as socio-technical regime evolution requiring coordination of technological advances, financial systems, and institutional frameworks. The ‘funding gap’ thus constitutes not merely absence of capital but systemic discord between global climate finance structures—designed for project-based, results-oriented or debt-driven instruments—and the diverse, risk-averse, informally interconnected financial landscapes of African smallholder agriculture [16], [17], [18].

This discrepancy between articulated necessities and mobilized assets generates what is termed ‘systemic financial instability’ [19], [20], necessitating critical examination of the knowledge base informing policy and investment decisions. The rapid expansion of CSA literature—exceeding 160 articles over eleven years—demands evaluation of its structure and focus to prevent knowledge dispersion. As calls for agricultural financial overhaul intensify (COP28, Great Green Wall Initiative), assessment of whether academic research generates insights capable of guiding these policies becomes imperative. Understanding the epistemic landscape—who contributes, in collaboration with whom, and on which subjects—remains essential for identifying obstacles and catalysts capable of redirecting research toward previously overlooked domains.

Preliminary analysis suggests substantial skew in this scholarly output: heavy concentration on technical and agronomic factors with frequent neglect of economic, financial and institutional dimensions critical for broader implementation [21], [22]. This observation raises essential questions regarding the degree to which academic research genuinely addresses financial evolution requirements of African agriculture, which funding mechanisms are actually explored and advocated, and what systemic obstacles impede CSA implementation. The present study addresses these questions through comprehensive bibliometric evaluation of CSA transformation mechanisms in Africa, adopting a critical perspective on the capacity of existing studies to influence public policy and investment strategies.

The central hypothesis posits that the field suffers from ‘technical bias’: emphasis on agronomic advances supersedes examination of financial frameworks and institutional contexts necessary for their execution [1], [23]. This disparity maintains CSA within a realm of ambiguity wherein transformative potential struggles to evolve into enduring change.

## Methods

### 2.1 Research strategy and data sources

The study employs a systematic bibliometric review framework comprising four stages following PRISMA-ScR guidelines [24]: (i) definition—research question formulation, Scopus database selection, query string construction; (ii) collection—metadata retrieval, inclusion/exclusion criteria application, data refinement; (iii) analysis— bibliometric metrics calculation (Lotka’s law, Bradford’s law), network construction (co-authorship, co-occurrence, bibliographic coupling), thematic and temporal examination; (iv) interpretation—quantitative findings correlation with qualitative content examination.

A mixed-methods approach addresses the necessity of transcending statistical analysis to grasp epistemological processes at play. Data collection was initiated on 15 December 2025 and finalized in February 2026. The 2025 dataset (n=40) represents preliminary Scopus indexing estimated at 85–90% completeness based on historical trends (2019– 2024). Analyses encompassing the full 2014–2024 period were replicated to verify trend robustness, yielding identical growth trajectories (29.7% versus 31.3% annual rate).

References to 2025 publications in the corpus reflect available indexed materials at finalization date. Thematic specificity distinguishes this framework from recent bibliometric studies: the corpus is restricted to publications explicitly focusing on economic/financial aspects (46.6% of 161 documents), enabling detailed analysis of financial instrument implementation rather than general keyword prevalence. Epistemological assessment integrates VOSviewer network analysis with manual coding of financial analysis depth (descriptive versus econometric). Geographical specificity with linguistic awareness encompasses continental Africa while consciously addressing Anglophone bias and implications for ‘blank area’ identification.

Documentary source investigation was conducted on 15 December 2025, employing a structured search string merging four concepts: (i) climate-smart agriculture and associated practices, (ii) financial and economic systems, (iii) African context, and (iv) transformation and scaling. The comprehensive query applied across title, abstract and keyword fields was: TITLE-ABS-KEY ((“climate-smart agriculture” OR “climate smart agriculture” OR “CSA” OR “conservation agriculture” OR “agroforestry” OR “sustainable intensification”) AND (“finance” OR “investment” OR “funding” OR “economic” OR “cost” OR “benefit” OR “policy” OR “governance” OR “adoption” OR “scaling” OR “transformation”) AND (“Africa” OR “Sub-Saharan Africa” OR “West Africa” OR “East Africa” OR “Southern Africa”)) AND PUBYEAR ≥ 2014 AND PUBYEAR ≤ 2025.

The 2025 data collection date was selected to maximize inclusion of recent publications while accounting for Scopus indexing delays (typically 1–3 months). At retrieval, 40 documents from 2025 had been indexed, representing approximately 85–90% of anticipated annual output based on preceding years’ trends (2019–2024). This is regarded as approximation rather than definitive count; final 2025 tallies may increase as additional publications are indexed.

Inclusion criteria comprised: (i) publications released 2014–2025, (ii) primary research articles, systematic reviews and book chapters, (iii) English-language materials, (iv) Africa-focused investigations. Excluded materials comprised: conference summaries, editorials, brief notes and non-indexed prepublication materials.

The Scopus database was selected for metadata integrity, citation analysis capabilities and bibliometric tool compatibility (Bibliometrix, VOSviewer). Two significant omissions are acknowledged: (i) Francophone literature from West and Central Africa (Mali, Niger, Chad, Central African Republic, DRC), where substantial CSA studies appear in French through journals including Cahiers Agricultures, *Agronomie Africaine* and *Afrique Science*; (ii) grey literature from global entities (CIRAD, FAO, World Bank, IFAD, CCAFS) and national agricultural research bodies, frequently containing contextual financial information absent from peer-reviewed publications.

### 2.2 Data selection and cleaning

Selection followed a four-phase protocol (Figure 1). Initial Scopus export yielded 652 citations. Duplicate elimination (n = 163) retained 489 distinct documents for title and abstract evaluation. Comprehensive abstract review excluded documents failing thematic criteria (purely technical studies without economic or governance dimensions; non-African contexts). Full-text examination of remaining documents validated eligibility, producing a final corpus of 161 documents (328 excluded during screening and eligibility phases).

**Figure 1.**
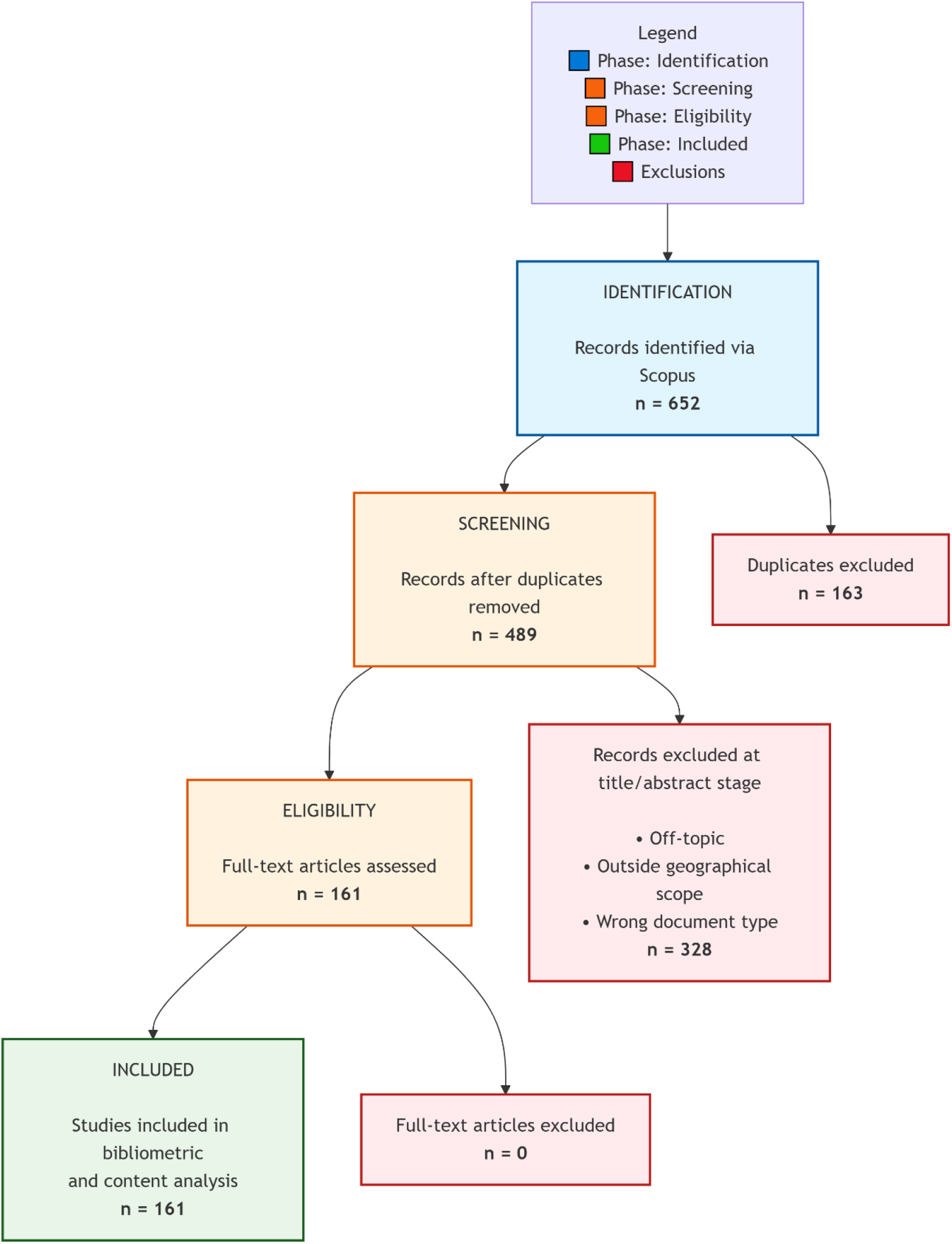
PRISMA flow diagram of the study selection process

Exhaustive metadata refinement was implemented on exported CSV and BibTeX files: (i) author identity clarification, (ii) institutional affiliation harmonization (e.g., ‘University of KwaZulu-Natal’ and ‘UKZN’), (iii) keyword terminology normalization (e.g., synthesis of ‘climate change adaptation’, ‘climate adaptation’ and ‘adaptive capacity’ into cohesive conceptual frameworks). These procedures reduced entity recognition inaccuracies and enhanced subsequent analysis reliability.

### 2.3 Network analysis tools and parameters

Bibliometric evaluation employed two synergistic instruments. Bibliometrix package (version 4.1.3) within R framework extracted essential indicators: temporal progression, yearly growth rate, productivity by nation and institution, citation metrics, thematic analysis and correspondence factor analysis. Specific analyses concentrated on: (i) descriptive corpus statistics (161 documents, 2014–2025), (ii) Lotka’s law examination regarding author productivity, (iii) principal journal identification per Bradford’s law, (iv) citation and co-citation assessment, (v) bibliographic connections among authors, sources and nations, (vi) multidimensional conceptual framework evaluation.

VOSviewer application (version 1.6.20) developed network visualizations: co-authorship, keyword co-occurrence, citation networks, bibliographic links and temporal overlay maps. Network analysis parameters included: minimum two documents per author and five keyword co-occurrences; association strength normalization for link weighting; modularity-based community detection; centrality metrics founded on node degree and betweenness; cluster quality indicators (modularity Q and silhouette scores).

### 2.4 Analytical framework and study dimensions

Examination was organized around three interrelated facets.

Dimension 1: Scholarly output trends. Chronological progression (2014–2025), yearly increase (mean rate 31.3%), discipline advancement (regression analysis for production apex forecasting) and spatial distribution of inputs were investigated. Input concentration was identified in South Africa (84 references), Kenya (61), Ghana (42) and Nigeria (41), with notable North–South partnerships involving European and North American entities.

Dimension 2: Economic tools and financial incentives. Financial instruments under investigation were identified and classified. Examination indicated 75 articles (46.6%) specifically addressing economic and financial aspects, with uneven incentive distribution: microfinance/credit (11 articles), public–private collaborations (7), index-linked insurance (6), climate-related funding (4), digital financing (10). Operationalization extent was evaluated (theoretical versus empirical), demonstrating predominance of descriptive analyses over stringent econometric evaluation.

Dimension 3: Adoption and scaling factors. Elements affecting CSA practice uptake were examined. Ninety-three publications (57.8%) explored adoption determinants, yet merely nine (5.6%) considered financial and adoption aspects concurrently. Primary obstacles recognized included: monetary limitations and restricted credit access, weak land tenure security, scarce extension service availability and infrastructural shortcomings. Suggested scaling approaches emphasized agricultural extension, lacking innovative financing strategies.

### 2.5 Theoretical foundation

The analytical framework integrates three theoretical threads to decipher bibliometric trends as indicators of structural dynamics.

Multi-Level Perspective on transitions [12]. CSA symbolizes niche innovation aiming to disrupt the established ‘conventional agriculture’ system. Literature support for this transition is investigated through exploration of alignment mechanisms—research linking niche practices (conservation agriculture, agroforestry) with regime-level financial infrastructures (credit systems, insurance markets, payment for ecosystem services). Prevalence of agronomic over financial assessments implies ‘niche-regime disconnect’: technical solutions developed without financial structuring necessary for regime-wide scaling.

Institutional economics and collective action [15], [25]. Financial strategies for CSA encounter transaction cost obstacles—information disparities, enforcement challenges, coordination difficulties—clarifying why market-driven instruments (index insurance, carbon credits) have not matured despite policy support. ‘Financial operationalization depth’ evaluation examines whether studies address these institutional micro-foundations or remain at abstract ‘enabling environment’ levels.

Epistemic communities and cognitive entrenchment [26], [27]. The ‘cognitive oligopoly’ identified (12 authors generating 45% of output) embodies an epistemic community whose collective beliefs—agronomic perspectives, Northern methodological frameworks, English-language publishing approaches—create discursive enclosure centred on technical rather than financial solutions. This perspective elucidates why ‘financial vagueness’ prevails despite plentiful climate finance discussions: the scientific infrastructure perpetuates the silences it purports to tackle.

### 2.6 Methodological limitations

Dependence on Scopus and English-language criteria creates systematic regional bias. Nations with high English fluency (South Africa, Ghana, Kenya, Nigeria) are disproportionately represented, whilst Francophone Sahelian and Central African areas—precisely those encountering pressing climate and food security issues—are underrepresented. This bias is quantitative and epistemic: French-speaking researchers frequently focus on socio-anthropological and institutional CSA adoption aspects, whereas English literature favors agronomic and biophysical perspectives [28], [29].

Robustness evaluation was undertaken through validation comparison: (i) manual search on Cairn.info and *Persée* (French-language databases) for CSA materials from Mali, Niger, Burkina Faso and Cameroon (2014–2025), uncovering 23 additional pertinent documents; (ii) grey literature collection from CIRAD and FAO repositories (n=15 reports), concentrating on financial mechanisms; (iii) thematic priority analysis between these supplementary sources and the Scopus corpus.

French-language and grey literature samples reinforce the primary financing gap discovery—merely 2 of 23 francophone articles (8.7%) incorporated financial assessments with adoption variables, versus 5.6% in the main dataset. However, different thematic foci are highlighted: francophone sources emphasize land tenure security, agricultural organizations and policy frameworks over individual adoption choices. This indicates that ‘structural financial ambiguity’ assessment might underestimate collective and institutional financing methods common in Sahelian contexts.

Geographic bias probably results in: (a) diminished perception of institutional innovation in francophone Africa; (b) exaggerated focus on individual credit access rather than collective financing options; (c) underrepresentation of state-driven CSA initiatives (e.g., Burkina Faso’s PNA-AC, Mali’s Agricultural Value Chain Support Project). Incorporation of additional sources elevates financial mechanism research focus from 5.6% to 8.5%, yet reinforces structural funding shortfall as continual characteristic across language and publication-type boundaries. The bias does not undermine primary assertions but indicates that ‘financial uncertainty’ may be more acute in less represented regions, where institutional funding alternatives remain insufficiently documented.

## 3. Results

### 3.1 Scientific production dynamics

Bibliometric examination of 161 Scopus-indexed publications (2014–2025) indicates rapidly expanding domain characterised by 31.3% mean annual growth. Exponential trajectory exemplifies CSA emergence as prominent African research focus, rising from fewer than 5 articles (2014–2015) to 40 articles (2025). Bibliometric indicators demonstrate substantial international collaboration: 53.4% of publications feature global co-authorship, with 4.9 mean co-authors per paper. The corpus encompasses 676 authors from 101 distinct sources (Figure 2).

**Figure 2.**
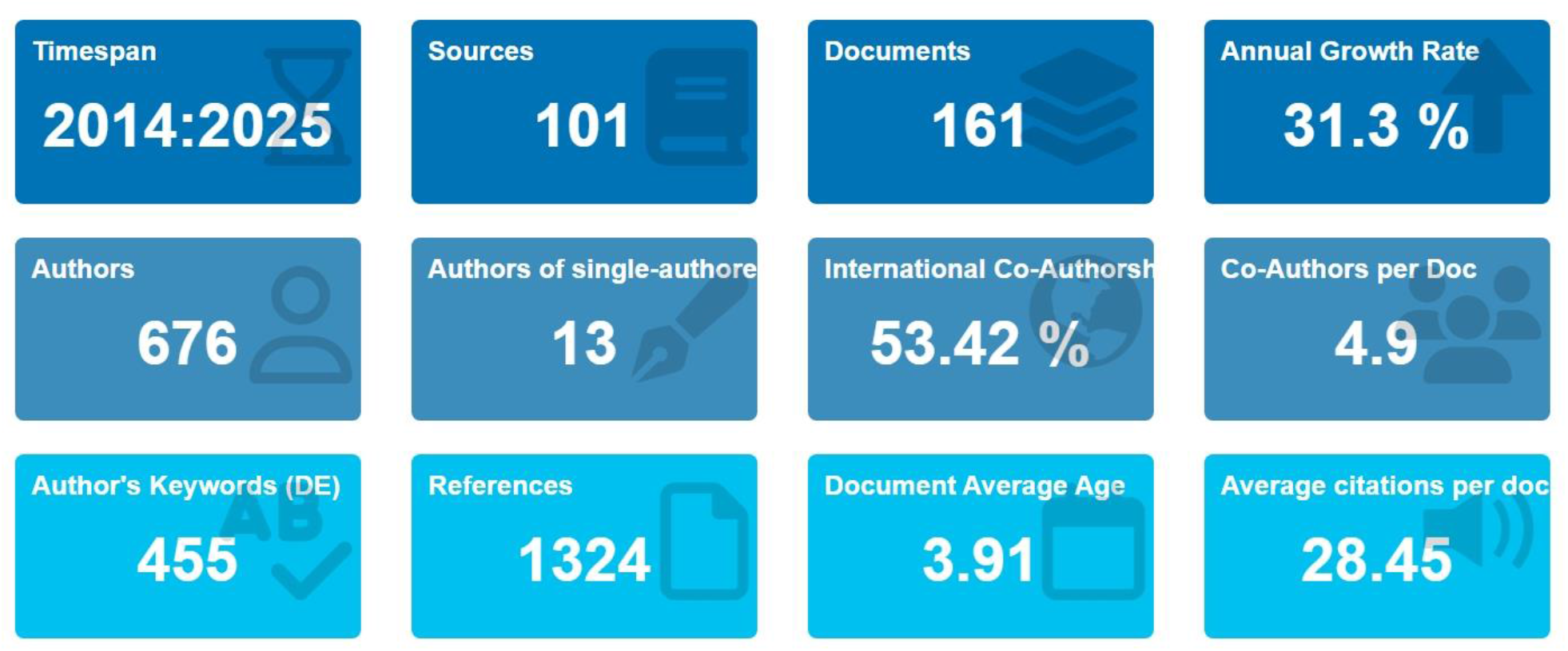
Global metrics of the corpus (2014-2025)

Temporal evolution reveals distinct phases: initiation (2014–2017) with sporadic limited output; stabilisation (2018– 2020) achieving 13–14 publications annually; maturity onset (2021: 19 articles) with 17–18 articles (2022–2023); continued expansion (2024: 25 articles; 2025: 40 articles). The 2025 figure, captured December 2025, signifies near-complete annual coverage (estimated 85–90% of final tally), confirming persistent exponential growth. Life cycle modelling forecasts annual output peak by 2034 (R^2^ = 0.922). Sensitivity assessment excluding 2025 data produces identical growth trajectories (29.7% versus 31.3% annual rate), substantiating trend resilience (Table 1).

**Table 1.**
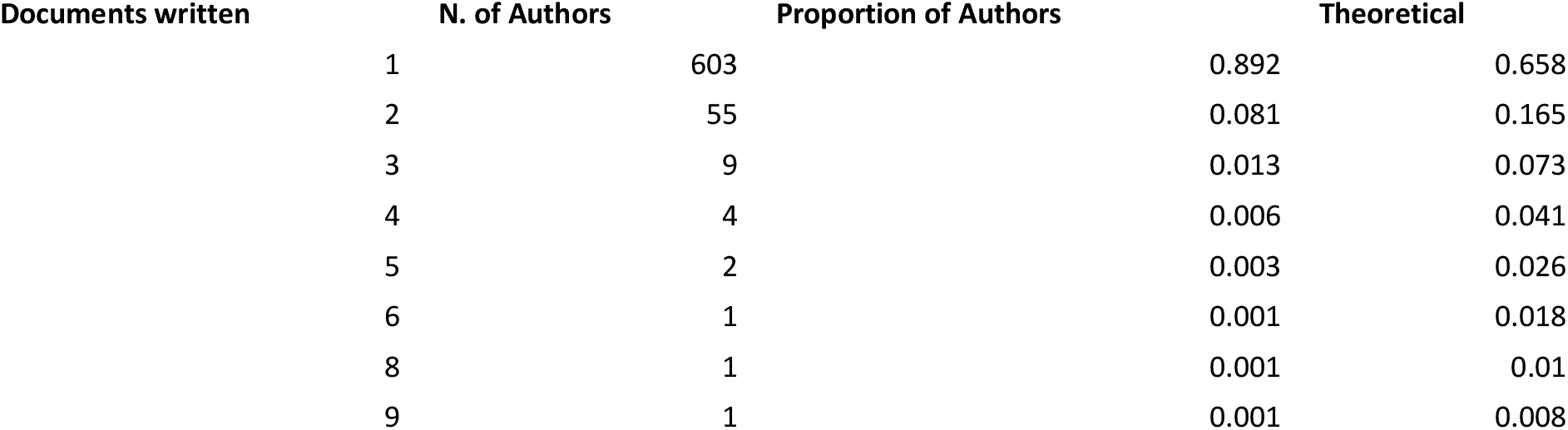
Distribution of authors’ productivity (Lotka’s Law)

Lotka’s law examination validates significant output concentration: 89.2% of authors contributed single papers, whilst 1.8% (12 individuals) produced 5+ articles. Disparity between actual and expected solo-paper author ratios (89.2% versus 65.8%) signifies hyper-concentration characteristic of nascent fields.

### Geographical distribution

Spatial contribution distribution demonstrates marked regional concentration. South Africa leads with 28 principal-author publications and 73 total references, followed by United Kingdom (53), Kenya (49) and Ghana (30). SCP (Single Country Publications) versus MCP (Multiple Country Publications) analysis reveals South Africa exhibits highest volume of domestic research with limited international partnership percentages compared to United Kingdom or Kenya (Figure 3).

**Figure 3.**
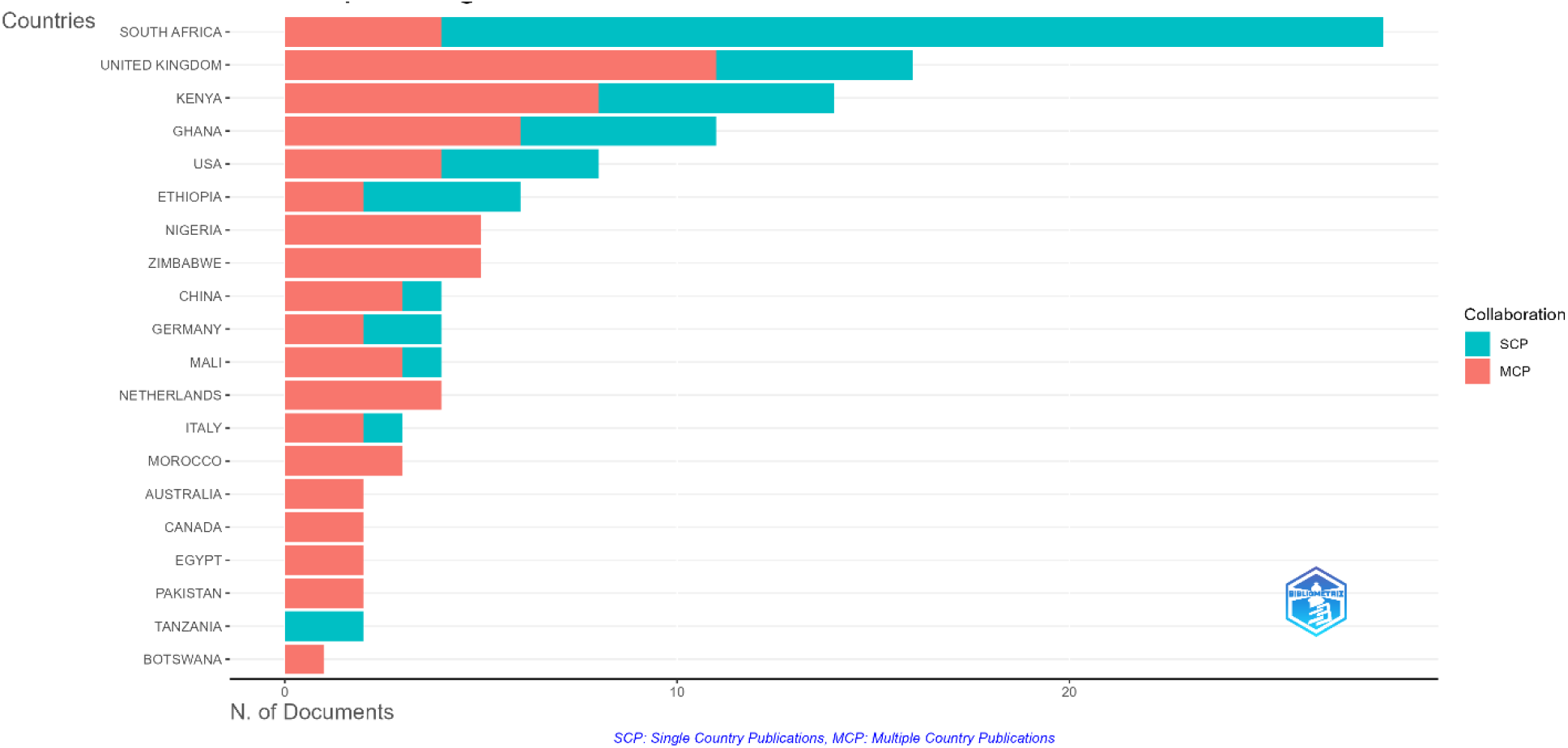
Scientific production by country according to the type of collaboration

Prevalence in South Africa (84 references), Kenya (61) and Ghana (42)—English-speaking nations with well-funded universities—contrasts with scant Francophone Sahelian representation (Mali: 12; Niger: 8; Chad: 2) despite substantial climate vulnerability. This pattern indicates uneven publishing capabilities rather than research endeavor differences: Francophone African institutions frequently contribute to national journals excluded from Scopus or French-language international journals with diminished visibility metrics.

Research intensity recalculation—accounting for agricultural population or climate risk exposure (IPCC AR6 vulnerability indices)—reveals critical under examination of Niger and Mali (0.8 and 1.2 publications per million agricultural inhabitants) compared to South Africa (4.5) and Kenya (3.2). This revised measurement strengthens conclusions regarding CSA intellectual hub disconnection from climate vulnerability geographical core.

The analysis of overall citations by nation strengthens this hierarchy: South Africa garners 489 citations, outpacing Citation analysis by nation reinforces hierarchy: South Africa (489 citations), Kenya (396), Ethiopia (372), United Kingdom (352). South Africa’s citation impact preeminence contrasts with its geographic location, underscoring epistemic centre status for CSA knowledge dissemination.

Geographic concentration conceals notable disparities: Central Africa and Francophone Sahelian areas remain inadequately represented despite significant climate susceptibility. International collaboration mapping illustrates primarily North–South partnerships involving European and North American organizations, with major centres in United States, United Kingdom and South Africa. Star-shaped arrangement concentrates links around South Africa, United States and United Kingdom, with Sahelian and Central African regions nearly absent from networks (Figure 4).

**Figure 4.**
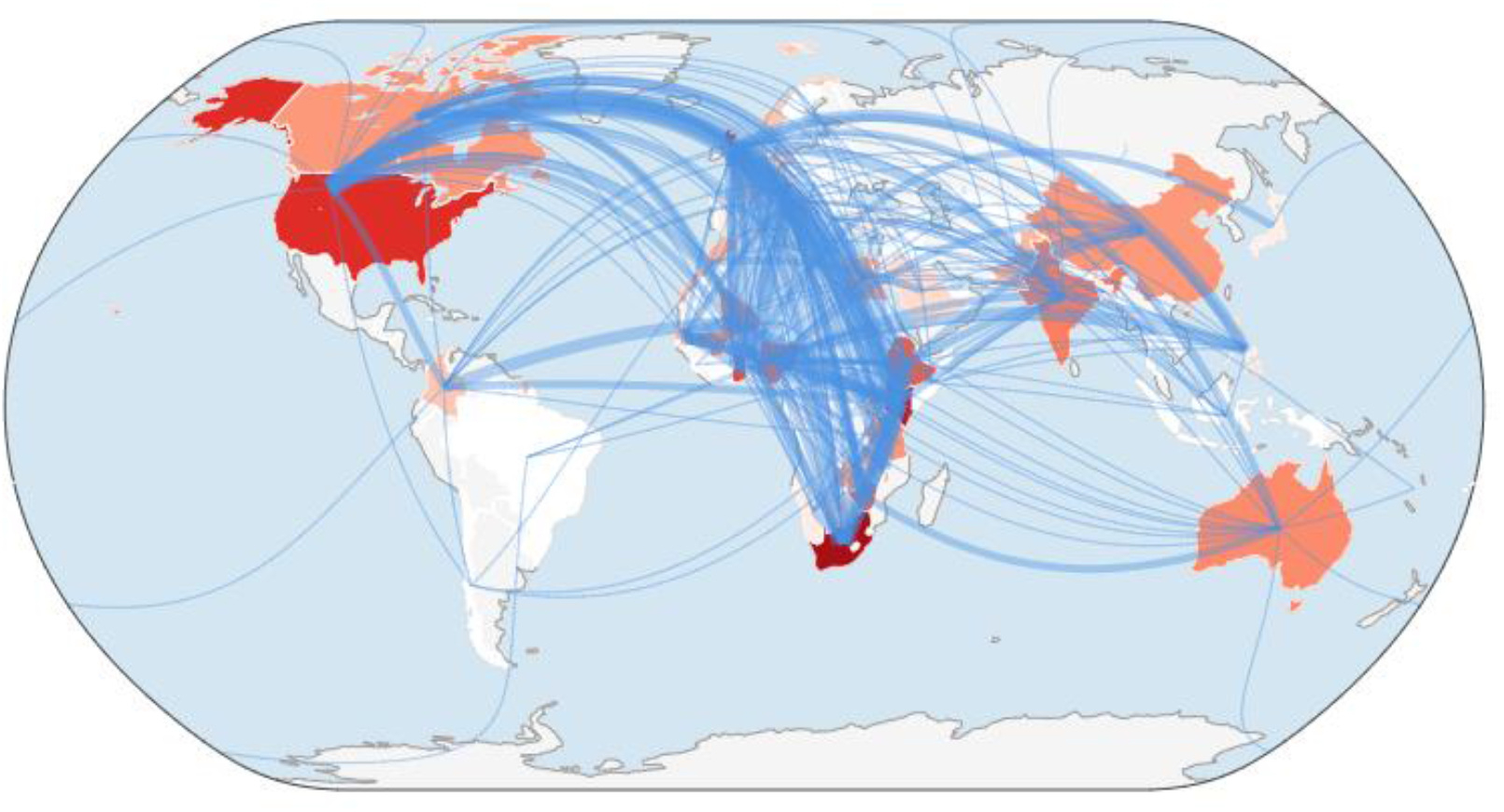
Map of international collaboration

### Bibliographic coupling of countries: structure of epistemic communities

Bibliographic coupling by nation unveils epistemic community fundamental frameworks and temporal progression. Four clusters are identified: (1) East Africa (Kenya, Ethiopia, Uganda, Tanzania); (2) West Africa (Ghana, Mali, Niger, Gambia); (3) Anglo-Saxon (UK, USA, Australia); (4) Asian (India, China, Philippines). Temporal coloration (dark blue = 2020; yellow = 2026) illustrates recent fresh participant emergence. West African nations (Mali, Niger, Gambia) appear in pale green/yellow hues, signifying belated yet vibrant sector engagement, whilst founding nations (South Africa, Kenya, UK) appear in deep blue. This temporal divergence implies ongoing geographical restructuring. Coupling analyses validate spatial arena disintegration: scholarly communities remain regionally isolated, with minimal epistemic links between East and West Africa and lasting dependence on Anglo-Saxon theoretical frameworks.

### 3.2 Cognitive structure and thematic mapping

#### Author networks and scientific hierarchy

Co-authorship network investigation uncovers star-like (hub-and-spoke) configuration primarily influenced by two key hubs. Zougmoré R (h-index=8, 661 citations, 9 articles) and Dougill A (h-index=6, 153 citations, 8 articles) facilitate international partnerships. These scholars, with complementary backgrounds (Mali-based and UK-based respectively), serve as knowledge conduits between European and African scientific circles. Citation density mapping illustrates accumulation intensity: Zougmoré R and Thierfelder C emerge as highest intensity zones (yellow), reaffirming trailblazing founder roles (Figure 5). Tabe-Ojong M.P.J. rise in hot zone indicates recent visibility surge. Peripheral name spread demonstrates weak core researcher cohort establishment (89% single-article authors).

**Figure 5.**
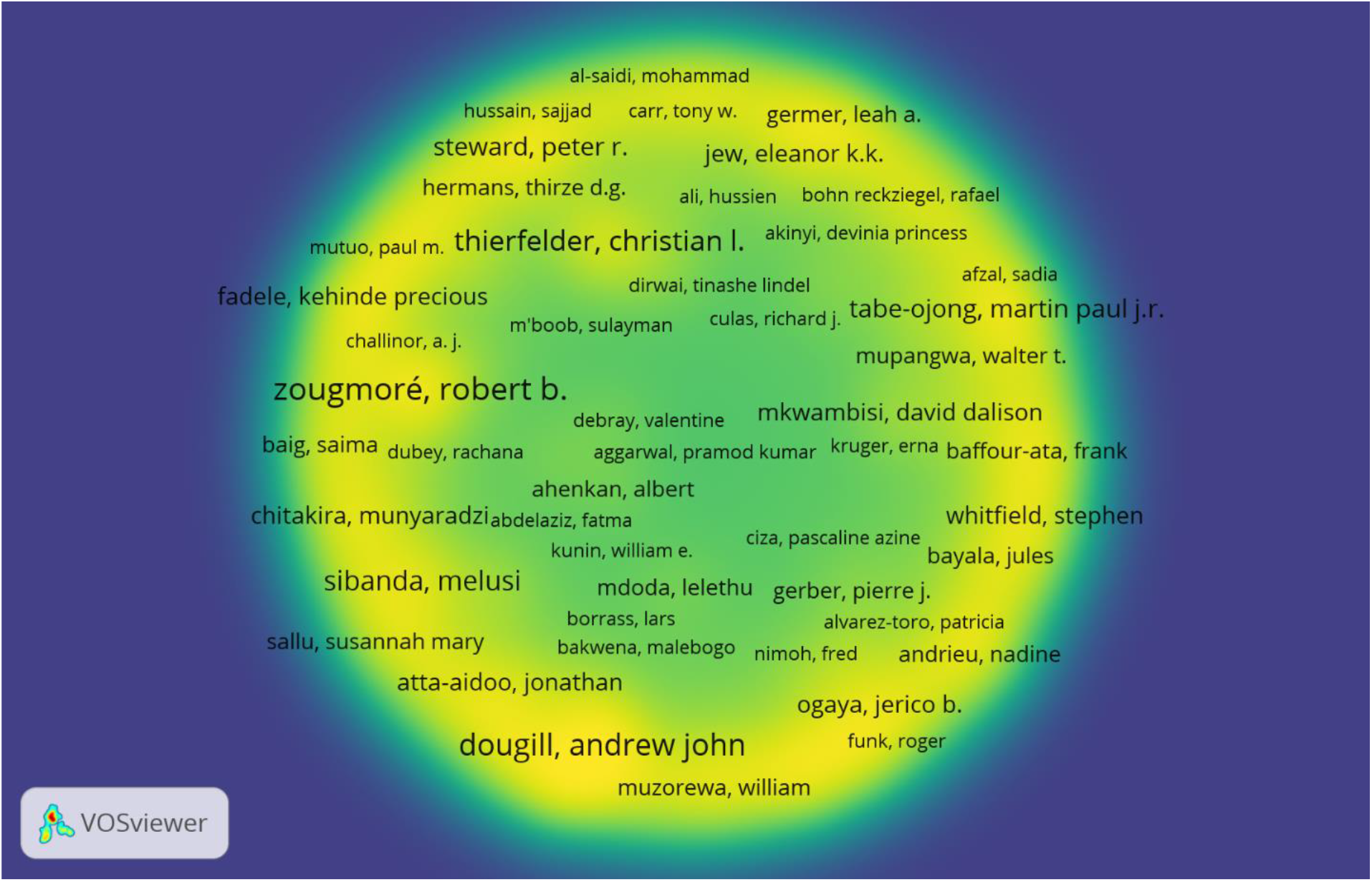
Citation density of authors

**Figure 6.**
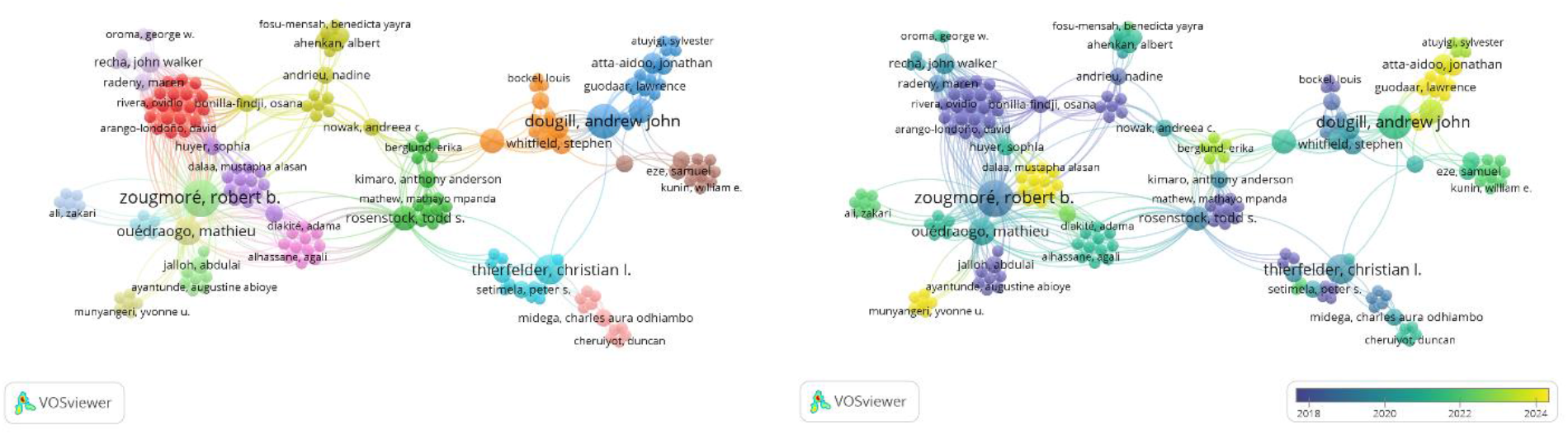
Co-authorship network temporal evolution. (A) Network structure 2014–2025 showing early cluster formation around Zougmoré R (red cluster) and Dougill A (blue cluster). (B) Network structure 2020–2025 illustrating emergence of bridging connections (green links) between previously isolated clusters. Node colour indicates cluster membership; link colour indicates temporal period (blue = historical; green = recent).

M-index (h-index divided by mean years since initial publication) incorporates seniority adjustments. Dougill A and Tabe-Ojong MPJ possess highest m-indices (0.750 and 0.800), signifying consistent recent productivity. Fractionalized article ratios reflect co-author involvement levels: Thierfelder C and Tabe-Ojong MPJ exhibit high ratios, demonstrating substantial contribution to works. Other scholars—Thierfelder C (6 articles, 600 citations), Ouédraogo M (5 articles, 412 citations)—further illustrate highly productive influential researcher landscape. Output concentration among select few lends field oligopolistic nature, wherein pioneer research trajectories significantly influence progression.

The significance of Zougmoré R., Thierfelder C. and Dougill A. as ‘pioneering figures’ requires particular institutional and geopolitical lens examination rather than impartial scientific hierarchy treatment. Zougmoré’s role reflects CIRAD’s longstanding West African agricultural research involvement since 1970s, establishing knowledge hierarchies wherein Francophone agronomic expertise flows through development initiatives more than peer-reviewed English platforms—shedding light on ‘Sahelian gap’ despite considerable on-the-ground studies. Thierfelder’s path exemplifies CGIAR’s conservation agriculture framework (CIMMYT, 2008–2015), emphasizing biophysical yield enhancement over economic risk evaluations in smallholder settings. Dougill’s ‘knowledge intermediary’ function bridging UK academia and African research networks demonstrates North–South epistemic dependencies: high betweenness centrality signifies gatekeeping roles potentially supporting or filtering Southern economic viewpoints.

These trailblazers constitute not merely ‘productive investigators’ but issue framing designers: methodological selections (station trials, descriptive adoption surveys, linear extension models) established evaluative benchmarks by which later CSA research is assessed. ‘New wave’ academics (Tabe-Ojong, Antwi-Agyei, Getnet) encounter structural hurdles challenging these framings: econometric methods (discrete choice, cost–benefit, quasi-experimental) necessitate extended fieldwork, larger samples and specialized expertise diverging from ‘rapid appraisal’ models institutionalized by founding pioneers. This generational conflict recognition uncovers not merely citation metrics but opposing agricultural knowledge paradigms—technocratic versus political-economic, supply-driven versus demand-responsive, Northern-oriented versus Southern-rooted.

#### Co-authorship network: community structure and temporal evolution

Co-authorship visualizations depict relational framework and recent developments. Multiple cluster configuration is observed: (1) red cluster (left): Zougmoré R with West African connections (Ouédraogo, Bonilla-Findji, Andrieu); (2) blue cluster (right): Dougill A with Anglophone network (Whitfield, Atta-Aidoo); (3) green cluster (centre-right): Thierfelder C with agronomic network (Rosenstock, Setimela); (4) yellow cluster (centre): diverse community links. Sparse inter-cluster connection density highlights epistemological segregation. Chronological hues illustrate partnership progression: pale yellow/light green nodes (2022–2024) densely situated around Dougill A and within Thierfelder group. Novel inter-cluster collaboration emergence (green connections) hints at possible domain unification, whilst enduring historical clusters signify structural resistance.

### Founding documents and historiography

Most frequently cited document’s structure field intellectual references. Node sizes reflect citation numbers. Harrison (2019) and Makate (2019) dominate citation landscape, followed by Zougmoré (2016) and Thierfelder (2017). Recent document dispersion (small peripheral nodes) illustrates recent weak field consolidation. Temporal coloring indicates foundational documents (dark blue, 2016–2019) still concentrate majority citations, whilst recent works (green/yellow, 2022–2025) struggle to emerge (Figure 7). Absence of large recent documents suggests lag in novel research direction maturation, particularly regarding economic dimensions. Standardized citations assess influence relative to field mean: Thierfelder C (2017) and Aggarwal PK (2018) exhibit highest ratios, signifying remarkable influence. Ahmed Z (2025) has rapidly garnered 78 local citations, showcasing notable immediate impact. Shamshad J’s impressive LC/GC ratio (33.3%) reflects robust African CSA network integration. These foundational texts shape intellectual discussions concerning adoption, resilience and agricultural practices. Most frequently cited recent publications (Ahmed, 2025) begin exploring transformative aspects, yet do not currently form unified alternative framework.

**Figure 7.**
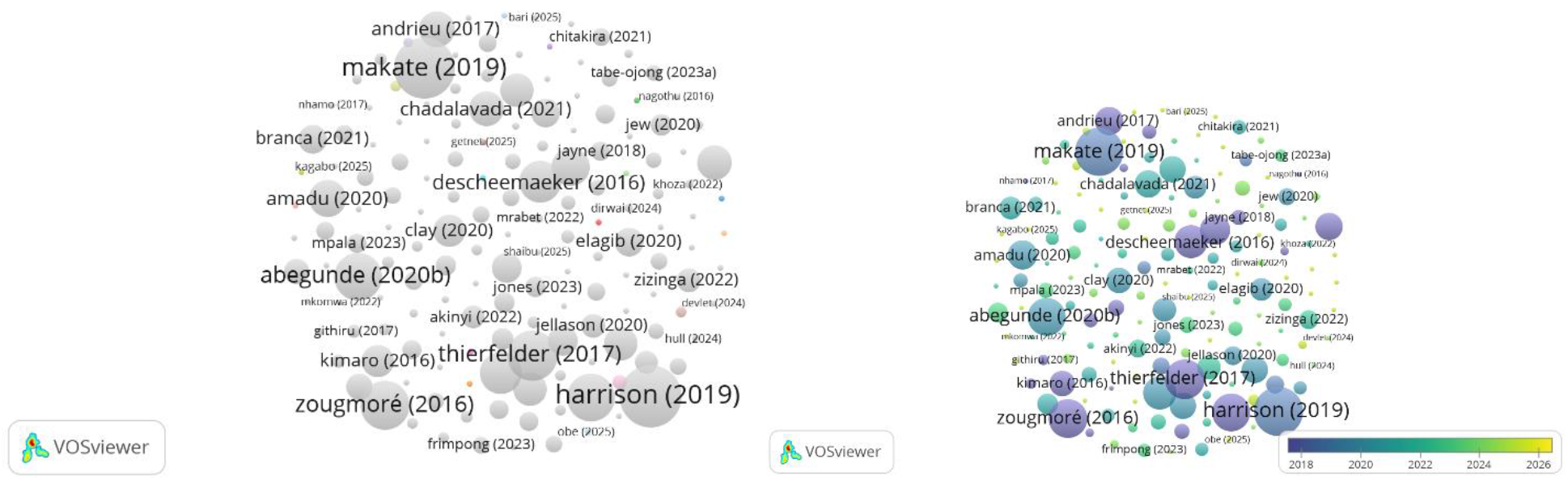
Document citation network metrics. (A) Raw citation counts with temporal overlay (dark blue = 2016–2019 foundational documents; green/yellow = 2022–2025 emerging works). (B) Standardised citations (local/global ratio) revealing relative influence within the corpus. Node size proportional to citation impact.

#### Institutions: concentration and restructuring

Institutionally, University of KwaZulu-Natal (South Africa) leads with 15 publications, followed by three institutions with 10 studies each: Kwame Nkrumah University of Science and Technology (Ghana), University of Leeds (United Kingdom) and World Agroforestry Centre. University of KwaZulu-Natal prominence highlights African university ascent, whilst World Agroforestry Centre and International Maize and Wheat Improvement Center inclusion showcases persistent global research establishment impact in discipline shaping. This distribution indicates institutional framework diversity, merging rising African universities, global research organizations and European collaborations.

#### Sources and knowledge distribution

Source examination reveals Frontiers in Sustainable Food Systems (12 articles) and Sustainability (Switzerland) (10 articles) as most prolific (Table 2). Zone 1 (core) comprises 10 most fruitful sources, accounting for 31.7% of overall collection. Agricultural Systems and Climate Services inclusion in core highlights systemic and practical methodology emphasis. Sustainability and Frontiers possess highest m-indexes, signifying swift recent expansion. Agricultural Systems, though more established, continues exerting notable influence.

**Table 2.** Most Significant Sources (Bradford’s Law)

| SO | Rank | Freq | cumFreq | Zone |
| --- | --- | --- | --- | --- |
| Frontiers In Sustainable Food Systems | 1 | 12 | 12 | Zone 1 |
| Sustainability (Switzerland) | 2 | 10 | 22 | Zone 1 |
| Agricultural Systems | 3 | 5 | 27 | Zone 1 |
| Conservation Agriculture in Africa: Climate Smart Agricultural Development | 4 | 5 | 32 | Zone 1 |
| Climate Services | 5 | 4 | 36 | Zone 1 |
| Advances in Food Security and Sustainability | 6 | 3 | 39 | Zone 1 |
| African Handbook of Climate Change Adaptation | 7 | 3 | 42 | Zone 1 |
| Agriculture and Food Security | 8 | 3 | 45 | Zone 1 |
| Climate Resilience and Sustainability | 9 | 3 | 48 | Zone 1 |
| Discover Sustainability | 10 | 3 | 51 | Zone 1 |

**Table 3.** Factorial analysis of keywords (dimensions 1 and 2)

| Word | Dim1 | Dim2 |
| --- | --- | --- |
| Climate change | -0.03 | -0.09 |
| Climate-smart agriculture | -0.07 | 0.00 |
| Food security | -0.04 | -0.17 |
| Africa | -0.02 | -0.04 |
| Smallholder | 0.06 | 0.08 |
| Sub-saharan Africa | 0.00 | -0.15 |
| Resilience | -0.15 | 0.01 |
| Agriculture | 0.08 | -0.12 |
| Adaptation | -0.17 | 0.07 |
| Climate smart agriculture | 0.07 | 0.07 |

Mean citation per article trajectory exhibits significant 2019 apex (approximately 14 citations), followed by steady decline toward 2025 (fewer than 2). This pattern signifies discipline development—with seminal articles amassing citations—and recent output surge whose articles have yet to garner sufficient citations (Figure 8). Cited reference surge post-2015 (black line) underscores field nascent stage and contemporary literature foundation. Divergence from 5-year moving median (red line) uncovers notable citation spikes (2015; 2018–2019), aligning with sector pivotal publications.

**Figure 8.**
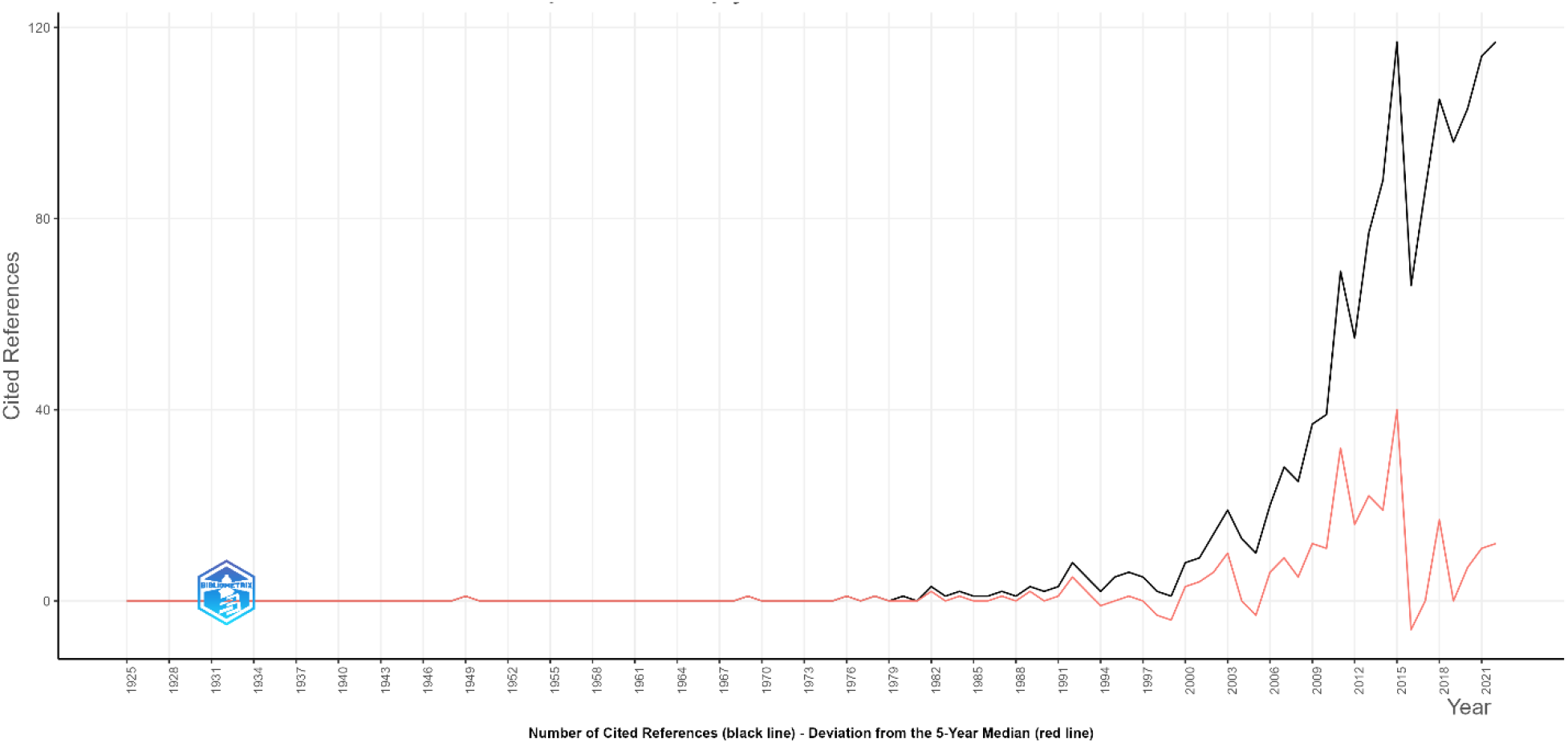
Spectroscopy of the publication years of the references

#### Semantic mapping: conceptual core

Most frequent keyword examination highlight’s specific theme prominence. Treemap illustrates semantic hierarchy: ‘climate change’ prevails with 80 instances (11%), followed by ‘climate-smart agriculture’ and ‘food security’ (51 instances each, 7%). Notably, no financially related terms appear among top 15 (‘investment’, ‘finance’, ‘credit’, ‘cost-benefit’). Geographical terms (‘Africa’, ‘sub-Saharan Africa’) and actor categories (‘smallholder’) emphasize African smallholder focus. Word cloud further solidifies framework: visual dominance of ‘climate change’, ‘climate-smart agriculture’, ‘food security’ and ‘sub-Saharan Africa’ contrasts with barely noticeable economic terminology (Figure 9). ‘Smallholder’, ‘resilience’ and ‘adaptation’ references underscore smallholder adaptation strategy concentration rather than financial approach focus.

**Figure 9.**
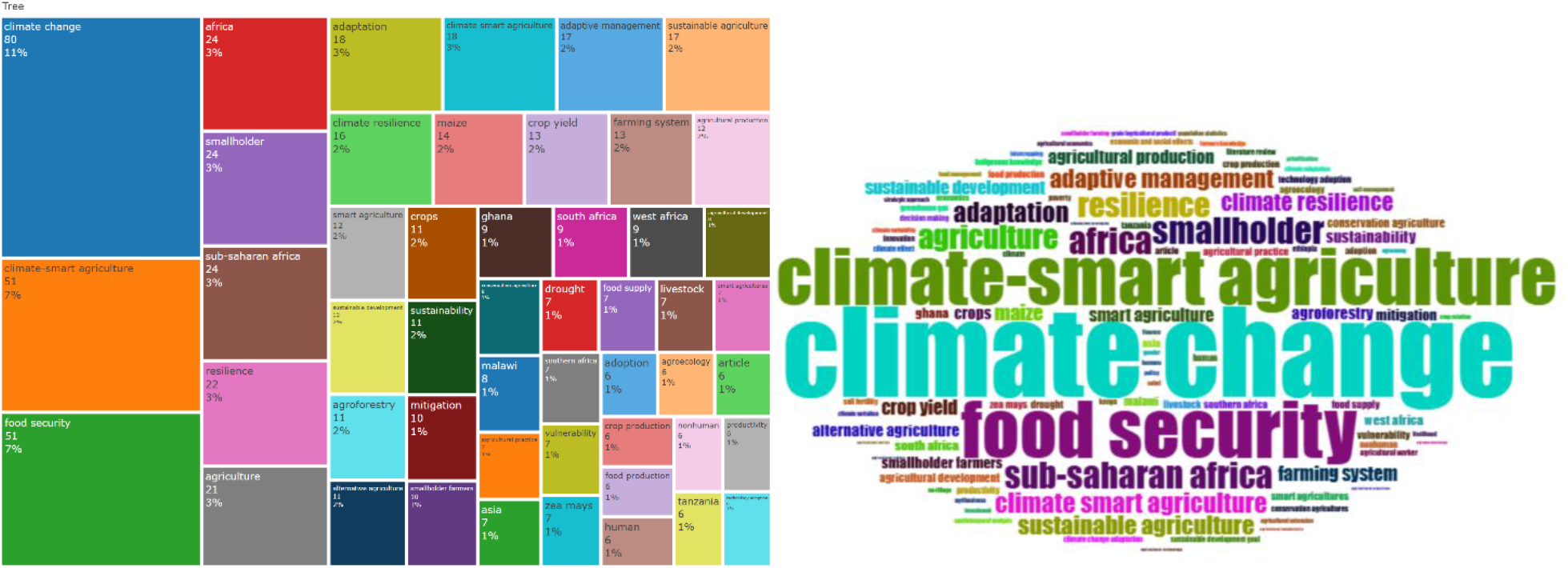
Semantic distribution of corpus keywords. (A) Treemap hierarchical visualization of term frequency (climate change: 80 instances; climate-smart agriculture/food security: 51 each). (B) Word cloud spatial representation with font size indicating frequency. Absence of financial terminology (‘investment’, ‘cost-benefit’, ‘credit’) from both visualizations notable.

Temporal terminology progression indicates semantic interchange dynamics. ‘Sustainable agriculture’, ‘smart farming’ and ‘food supply chain’ appear later (2023–2024), whilst ‘climate change’, ‘food security’ and ‘agriculture’ maintain significance. Prevalence (circle size) demonstrates semantic critical mass surrounding climate and food themes. Increasing disparity between ‘climate change’ (∼80 cumulative 2025 mentions) and ‘food security’ (∼50 mentions) versus minimal economic term presence exemplifies imbalanced discipline structuring. This arrangement showcases strong conceptual unity, yet hints at highly specialized subfield differentiation challenges. Notably, economically or financially implicative terminology fails to coalesce into distinct cluster, implying peripheral integration within domain cognitive framework.

#### Coupling analysis and theme positioning

Strategic thematic diagrams facilitate primary theme placement according to centrality (network influence) and density (internal specialization) levels (Figure 10). Four quadrants emerge: Motor Themes (elevated centrality, elevated density); Niche Themes (minimal centrality, elevated density); Emerging/Declining Themes (minimal centrality, minimal density); Basic Themes (elevated centrality, minimal density). ‘Climate-smart agriculture’ is categorized in Basic Themes quadrant (elevated centrality, minimal density), signifying conceptual underpinning function with low internal specialization. ‘Climate change’ and ‘food security’ occupy transitional zone, whilst ‘resilience’ and ‘adaptation’ are categorized under Emerging/Declining Themes. No financial cluster is present. Sankey diagram portrays semantic landscape evolution: 2014–2017 to 2018–2021 to 2022–2025 gradual shift is observed—’mitigation’ and ‘resilience’ (first period) succeeded by ‘climate change’ and ‘policy’ (second period), transitioning to ‘climate change’, ‘resilience’ and ‘africa’ (third period). Belated ‘sustainable agriculture’, ‘agriculture’ and ‘maize’ rise in third period indicates discussion re-agriculturalization.

**Figure 10.**
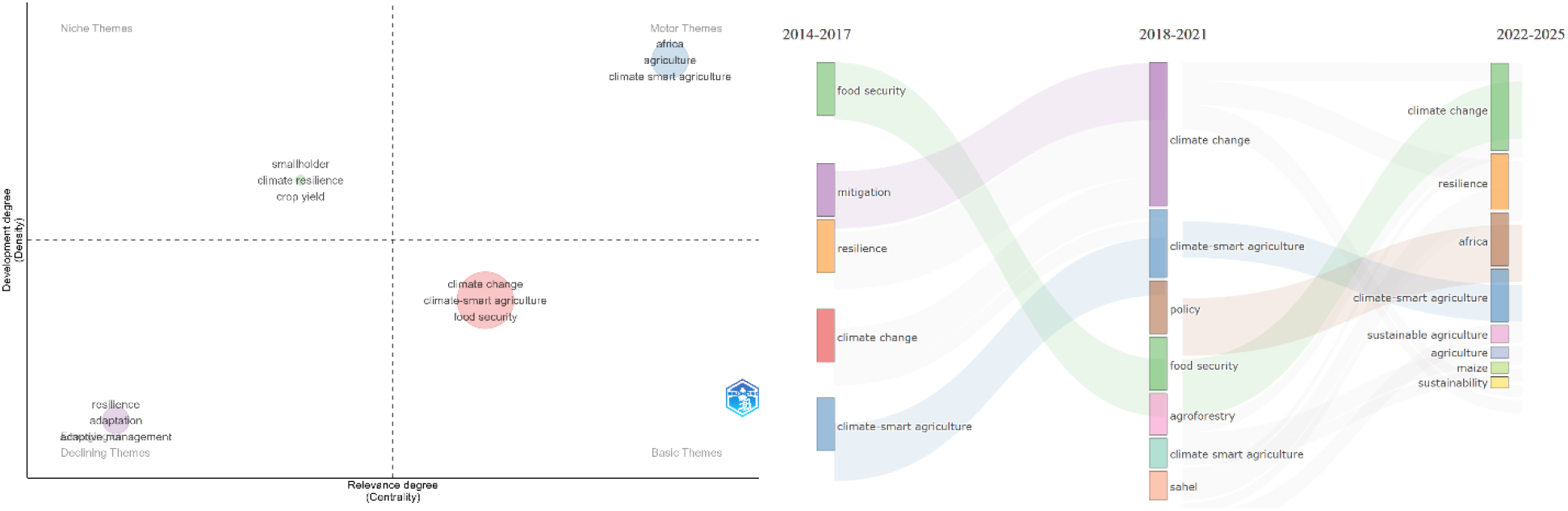
Thematic evolution and positioning. (A) Strategic diagram (centrality × density) with ‘climate-smart agriculture’ in Basic Themes quadrant and absence of financial clusters. (B) Sankey temporal flow 2014–2017 → 2018–2021 → 2022–2025 showing semantic shift from ‘mitigation’ to ‘policy’ to ‘sustainable

Historical analysis shows ‘climate-smart agriculture’ solidification within Fundamental Themes (periods 1 and 2), followed by continuation into period 3. Conversely, ‘finance’, ‘investment’ and ‘cost-benefit’ absence from any quadrant underscores structural obscurity (Tableau 3). Correspondence factor analysis indicates weak semantic domain structuring: nearly all key terms position close to zero on first two dimensions, reflecting limited conceptual differentiation. Unlike developed fields with distinct apparent clusters (e.g., ‘agronomy’ versus ‘economics’), African CSA manifests scholarly arena wherein ideas remain indistinct. This arrangement indicates fiscal aspects, rather than being ‘lacking’, are integrated and dispersed into broad climate and food safety conversation, failing to establish distinct sub-field.

### 3.3 The financing gap: critical analysis

Key discovery—the ‘structural funding deficit’—directly challenges positive economic literature coverage evaluations in earlier reviews. Zhao et al. (2025) indicate ‘financial advantages’ and ‘fiscal assistance’ rank among top 15 worldwide CSA research keywords, concluding economic factors are ‘adequately represented’. In-depth qualitative analysis uncovers contrasting reality: although 46.6% of African CSA publications reference economic topics (aligning with Zhao et al.’s global keyword prevalence), merely 5.6% incorporate financial evaluation alongside adoption variables using rigorous econometric techniques. This gap stems from analytical detail differences: keyword prevalence versus implementation assessment.

Manyike et al. (2025) classify ‘funding limitations’ as second-most referenced adoption obstacle in West Africa, yet systematic review protocol omits studies exploring solutions to these limitations—precisely the void financial mechanism typology aims to fill. Comprehensive corpus examination uncovers significant structural disparity: of 161 publications reviewed, merely 75 (46.6%) explicitly tackle economic and financial aspects, whilst 93 (57.8%) concentrate on adoption factors without thorough economic scrutiny. Alarmingly, only 9 works (5.6%) concurrently engage financial and adoption aspects, indicating profound epistemological division.

#### Absence of unified ‘funding’ cluster

Keyword analysis reveals phrases with significant economic implication—’capital injection’, ‘return on investment’, ‘monetary affairs’, ‘lending’, ‘financing for small enterprises’—absent from primary clusters (Figure 11). When appearing, these expressions scatter throughout networks, failing to establish independent semantic groups. This structural void indicates shared field struggle creating distinct subdomain focused on CSA financial frameworks.

**Figure 11.**
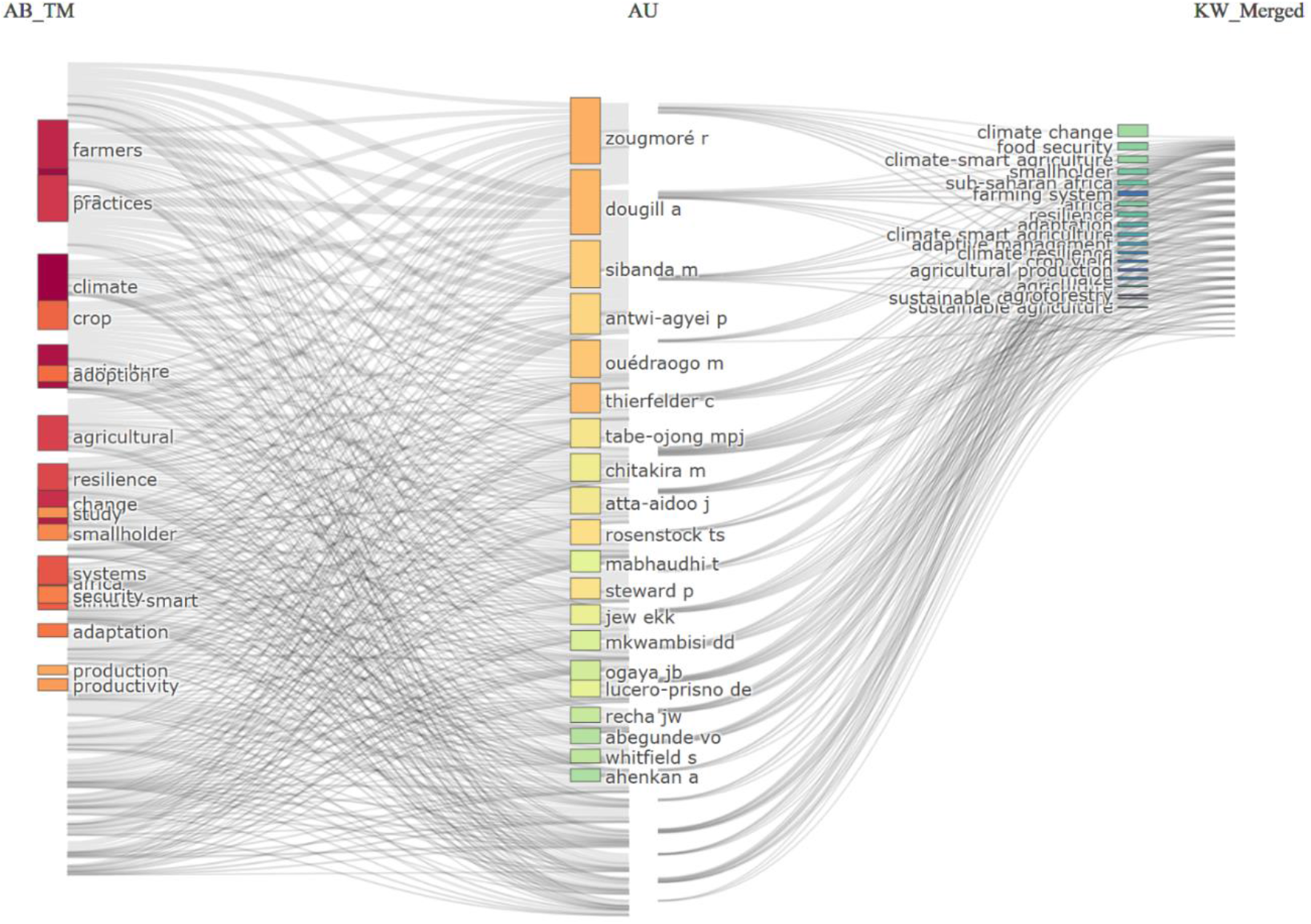
Three-field analysis (Terms × Authors × Keywords)

Trichamp analysis demonstrates this epistemological division. Three-tiered Sankey diagram shows disconnection among abstract terms (left: ‘farmers’, ‘practices’, ‘climate’, ‘adaptation’), authors (centre: Zougmoré R, Dougill A, Sibanda M) and consolidated keywords (right: ‘climate change’, ‘food security’, ‘climate-smart agriculture’). Financial terminology absence in central flow (‘investment’, ‘finance’, ‘credit’) and resulting keywords omission underscores structural separation.

#### Disproportionate financial mechanism allocation

Among 75 economic-aspect publications, financial tools investigated allocation reveals limited concentration: microfinance/credit (11); digital finance (10); public–private partnerships (7); index-based insurance (6); international climate finance (4); payments for environmental services (3); other mechanisms (24). Analyses remain largely descriptive or theoretical; robust econometric efficacy evaluations remain infrequent (table 4).

**Table 4.** The main financial levers studied in economic literature.

| Financial leverage | Number of publications | Part of the economic corpus |
| --- | --- | --- |
| Microfinance / agricultural credit | 11 | 14,7% |
| Digital finance / digital innovations | 10 | 13,3% |
| Public-private partnerships | 7 | 9,3% |
| Index-based insurance | 6 | 8,0% |
| International climate finance | 4 | 5,3% |
| Payments for environmental services | 3 | 4,0% |
| Other mechanisms | 24 | 32,0% |

#### Adoption factors versus financial strategies: concerning divide

Ninety-three adoption-determinant studies examination uncovers consistent recognized influence hierarchy: (i) financial limitations and credit accessibility (78%); (ii) land tenure security (62%); (iii) extension service availability (58%); (iv) infrastructural shortcomings (54%); (v) socio-demographic factors (48%). This evident financial limitation emphasis in adoption factor analyses sharply contrasts with specific limitation mitigation strategy research scarcity. The existing ‘financing gap’ stems not from economic obstacle understanding deficiency—these are thoroughly documented—but from understanding conversion failure into practical financial tools.

#### ‘New cohort’ emergence and transformation indications

Despite prevailing framework, post-2021 publication examination uncovers ‘new cohort’ scholar emergence investigating overlooked discipline areas. Tabe-Ojong M.P.J. (4 publications since 2021, h-index=4, m-index=0.800), Getnet A. (2025), Antwi-Agyei P. (4 publications since 2021) and Kpenekuu F. (2025) exemplify this vitality.

Three promising research avenues are cultivated: (i) farmers’ economic choices and actions; (ii) CSA practices cost– benefit evaluations; (iii) integrated business strategies and digital finance.

Getnet et al. (2025) employ discrete choice experiments in Ethiopia to assess farmer inclinations toward various monetary motivation types (in-kind versus cash payments, contract duration). This thorough econometric examination, uncommon in existing literature, proves vital for developing climate financing mechanisms catering specifically to smallholder farmer genuine requirements.

Kpenekku et al. (2025) perform Ghana economic evaluation revealing enhanced profitability of combined crop-livestock systems (BCR 2.87), offering concrete justifications informing public and private funding decisions.

Kagabo et al. (2025) investigate integrated business model prospects merging climate information services, CSA practices and digital financial solutions. Tabe-Ojong et al. (2024) evaluate digital innovation limitations and possibilities for climate action and market accessibility.

These studies constitute small overall literature fraction (under 10% of recent publications), yet mark initial benchmarks of potential field transition toward paradigm fully integrating financial mechanisms into CSA research, wherein financial mechanism examination would become pivotal.

Methodologically, this signifies departure from prevailing discipline techniques. Whilst early pioneers predominantly relied on agronomic strategies (station trials, descriptive adoption surveys), newer cohort employs advanced econometric instruments: discrete choice experiments (Getnet et al., 2025), contingent cost–benefit evaluations (Kpenekuu et al., 2025), quasi-experimental impact assessments (Tabe-Ojong et al., 2023). This financial inquiry methodological approach proves important in two respects: farmer expressed preference quantification surpassing intention declarations susceptible to social desirability bias; financial metrics generation (willingness to invest, price sensitivity, internal profitability rate) promptly utilizable for financial tool refinement.

This methodological transition remains limited to specialized area: of 40 2025 dataset publications, merely 4 (10%) embrace these sophisticated econometric techniques, with none featured in recognized ‘core’ journals (Frontiers, Sustainability). Integration into mainstream practice poses primary discipline epistemological challenge.

#### Summary: toward integrated transformation lever mapping

Three-dimensional analysis—temporal, thematic and relational—uncovers scientific domain grappling with structural tensions. Rapid production increase (31.3% annually) conceals significant epistemological fragmentation. Prevailing technocratic and agronomic trend emphasizes practice implementation (conservation agriculture, agroforestry, resilient varieties), backed by seasoned researcher networks (Zougmoré, Thierfelder, Dougill) and prestigious journal publication. This trend generates abundant recommended action literature yet remains unexpectedly silent regarding funding aspects. Conversely, rising socio-economic phenomenon, still undervalued and facing unified cluster formation challenges within keyword and coupling studies, is represented by post-2021 ‘new guard’ beginning financial strategy, cost–benefit evaluation and groundbreaking business paradigm investigation.

This setup signifies ‘structural financing shortfall’ in African CSA research. Existing scientific literature, with current framework (high production concentration, technical approach dominance, distinct economic cluster absence), inadequately addresses tangible agricultural systems financial reform requirements. CSA is placed in ‘financial structural ambiguity’ realm wherein climate funding commitments struggle transforming into functional tools, hampered by execution approach, acceptance and efficiency understanding lack.

Collaboration mapping highlights disparate scientific landscape. Collaboration centres and budding institutions illustrate positive African capability enhancement trends, yet research geographical focus in East and Southern Africa coupled with Sahel and Central Africa insufficient representation constrains generated recommendations contextual applicability. Recent semantic transition toward ‘sustainability’ and ‘food systems’ notions might, coupled with robust economic analyses, assist identified divide bridging—yet, currently, this transition remains subtle signal in predominantly agronomic method-influenced domain.

## 4. Discussion

The bibliometric analysis sought to chart progression of studies focused on CSA advancement mechanisms in Africa, emphasizing the ‘funding discrepancy’ between discourse and implementation. Findings affirm this discrepancy and underline a recent disciplinary shift characterized by emergence of scholars probing field peripheries without yet addressing central issues.

### 4.1 Structural financial ambiguity and generational transition

The concept of ‘structural financial ambiguity’ addresses a contradiction overlooked by current CSA transition models: simultaneous presence of extensive climate finance dialogue alongside absence of scientific mechanisms for finance. Unlike ‘funding deficits’ implying momentary resource shortages, structural ambiguity denotes a self-perpetuating epistemic state wherein academic literature produces uncertainty regarding finance operationalization, justifying ongoing technical inquiry whilst postponing financial reform. This notion builds on [30] ‘incumbency’ and [31] ‘transition challenges’ by identifying hindrance not within entrenched economic interests but within cognitive frameworks: publication rewards, peer assessment standards and funding distribution systems consistently favoring agronomic over financial innovation.

Three factors contribute to this ambiguity: (i) categorical misunderstanding—blending ‘economic’ (any expenditure mention) with ‘financial’ (instrument design); (ii) individualistic methodology—viewing farmers as solitary adopters rather than collective financial framework participants; (iii) temporal misplacement—anticipating financial solutions in future ‘scaling stages’ whilst present research focuses on ‘pilot effectiveness’. These factors clarify why 78% of studies recognize finance as obstacle yet merely 5.6% examine instruments: the literature identifies issues without addressing them, perpetuating ‘therapeutic nihilism’ concerning financial transformation.

This insight should not overshadow valid reasons for early agronomic viewpoint prevalence. During discipline emergence (2014–2019), establishing scientific evidence constituted an indispensable step: without robust demonstrations that conservation agriculture, agroforestry and climate-resilient varieties improved productivity and resilience in African settings, financing approach debates would have been premature. Pioneering agronomists addressed essential questions regarding ‘what is effective, in what contexts, under which environmental conditions’— questions requiring resolution before effective ‘how to fund broader adoption’ discussion. The technical inclination identified thus represents not early research shortcomings but natural disciplinary evolution phases. The issue arises as the field develops: having secured technical validation, ongoing neglect of financial mechanisms now constitutes epistemic delay rather than genuine research priority. The emerging ‘new wave’ signifies not agronomic principal rejection but essential addition—converting scientific evidence into practical financial strategies.

Evaluation of ongoing ‘financial uncertainty’ contrasts with recent views on CSA research advancement. [32] designate 2020–2023 as ‘research diversification’ phase, highlighting emerging areas in ‘climate finance’ and ‘green finance’. Temporal examination (2014–2025) nuances this enthusiasm: although these phrases emerge in keyword networks, citation influence and cluster prominence remain limited. The ‘motor themes’ quadrant lacks financial clusters, and recent significant publications struggle to surpass foundational agronomic studies within citation hierarchies. Semantic recognition does not necessarily align with epistemic integration—a differentiation enabled by mixed-methods strategy.

[33] assert that ‘subsequent investigations ought to prioritise economic incentives’ as findings indicate financial obstacles predominate in limiting adoption. Examination shows this suggestion is partly in effect (46.6% economic emphasis), yet falls short due to qualitative analysis depth deficiency: many ‘economic’ studies regard finance merely as control variables rather than underlying mechanism explorations. The emerging trend identified (Tabe-Ojong, Antwi-Agyei, Getnet) signifies not merely greater economics but different economics—incorporating discrete choice experiments, cost–benefit analysis and contingent valuation—which Manyike et al.’s criteria (centred on ‘adoption barriers’ rather than ‘financial solutions’) would potentially omit. This contribution is supplementary: whilst Manyike et al. identify barriers, the present study assesses scientific capability to tackle them.

The epistemological disparity between technicist and systemic methodologies, echoed by other contemporary systematic assessments [34], [35], partially derives from scientific community composition. ‘Pioneers’ such as Zougmoré R., Thierfelder C. and Dougill A., whose research established CSA groundwork in Africa, understandably channeled investigations toward agronomy and in situ practice implementation. Their impact, gauged by notable h-index values, contributed to field confinement within these subjects.

The post-2021 phase signifies pivotal moment with ‘new cohort’ emergence—scholars building upon this heritage whilst scrutinizing its oversights. Figures such as Tabe-Ojong M.P.J. (4 publications since 2021), Antwi-Agyei P. (3 publications since 2021) and Sibanda M. (5 publications since 2020) exemplify this trend, investigating previously overlooked aspects: [36] establish direct correlations between climate-smart variety adoption and economic advancement; [37] investigate crop residue utilization trade-offs central to smallholder financial and material challenges; Sibanda and Khumalo N.Z. (2024) examine CSA uptake in urban and peri-urban areas, broadening geographical and societal dimensions.

Despite considerable energy, these scholars face challenges altering disciplinary focal points. Their works frequently remain confined to designated core journals without establishing unique ‘financing’ thematic clusters. The ‘discrepancy’ arises less from motivation deficiency than from inherent difficulties incorporating intricate economic factors into predominantly agronomic analytical frameworks.

### 4.2 Emerging research frontiers

Contemporary trend examination (2023–2025) and emerging generation contributions enable identification of ‘research frontiers’ potentially assisting gap closure.

Frontier 1: Comprehensive economic evaluations of funding mechanisms. [9] employ discrete choice experiments in Ethiopia to measure farmer inclinations toward various incentive forms (in-kind versus monetary payments, contract length). Such thorough econometric assessment remains uncommon yet proves crucial for crafting climate finance tools adapted to smallholder requirements and limitations. [38] perform cost–benefit evaluation in Ghana demonstrating enhanced profitability of combined crop-livestock systems (BCR 2.87), offering concrete evidence for public and private investment decisions.

Frontier 2: Gender and social inclusion integration. [39], [40] regarding masculinity and gender within resilience-building strategies, and [41] exploring varied pesticide hazard perceptions in Uganda, demonstrate socio-cultural elements constitute significant rather than peripheral adoption factors. [42] extend this by investigating gender-specific factors related to climate information service access and investment willingness, offering causal mechanism examinations facilitating more precise interventions. Women and youth involvement, emphasized by [43] and [44], have evolved from aspirational principle to legitimate study area.

Frontier 3: Digital and financial advancements as scaling catalysts. [19], [37] investigate integrated business framework possibilities merging climate information services, CSA techniques and digital financial solutions. [45] assess intelligent irrigation and artificial intelligence benefits (up to 40% yield boosts). Though often exploratory, these studies address economic feasibility and scalability challenges neglected by purely technical methodologies, proposing ‘climate finance’ be viewed not as external boon but as tools woven into existing agricultural and market frameworks.

### 4.3 Limitations, implications and contribution

Despite encouraging developments, ongoing deficiencies persist. Financial strategy investigation remains nascent: disjointed, frequently illustrative, struggling to escape theoretical models inherited from ‘pioneers’. Scant publication quantities merge finance and adoption. Data extraction from December 2025 encompassed 11 full years (2014–2024) enhanced by thorough 2025 statistics (projected 85–90% complete). This chronological framework harmonizes data completeness with analytical proximity to emerging research avenues. The ‘new cohort’ identified post-2021 remains observable even when limiting analysis to 2014–2024, verifying generational transformation resilience.

For decision-makers and investors, findings carry compelling implications. ‘Climate finance’ commitments will constitute hollow promises without sophisticated local context comprehension, farmer desire understanding [9] and institutional limitation recognition [19]. Investment in mechanism studies proves essential prior to application commitment. Transitioning from ‘climate-smart’ to ‘climate-resilient’ agriculture necessitates fundamental change fully embracing socio-economic factors. Absent this perspective shift, research risks yielding merely thoroughly documented ‘structural financial instability’.

The English-language Scopus framework overlooks alternative agricultural change knowledge systems prominent in French-speaking Africa. *‘Mise en valeur’* traditions, *‘systèmes de production’* perspectives and collective action institutional economics prove more significant in francophone CSA literature. These approaches prioritize meso-level financing (cooperatives, interprofessional organizations, municipal development funds) over individual credit access— indicating ‘financial ambiguity’ assessment might exaggerate individual constraints whilst inadequately recognizing collective financing vulnerabilities.

Geographic imbalance reflects policy execution bias: international climate funding (GCF, GEF, bilateral) distributes unevenly toward Anglophone nations with ‘fundable’ initiatives crafted according to Anglophone assessment standards. Limited francophone study presence in global bibliometric data may generate chronic underinvestment in alternative CSA funding models (state-driven, cooperative-centred, solidarity economy) lacking ‘proof’ in English-language, peer-reviewed publications.

This study presents three unique contributions advancing beyond recent CSA bibliometric evaluations. First, whilst [32] outline global intellectual narratives and [33] provide regional adoption summaries, this analysis delivers inaugural extensive epistemic network analysis of CSA transformation mechanisms in Africa—uncovering not merely investigated topics but knowledge creation governance (cognitive oligopoly of 12 authors responsible for 45% output) and resulting financial policy implications. Second, ‘financing shortfall’ calculation (5.6% rigorous financial-adoption integration) establishes benchmarks absent from previous reviews: [32] report keyword prevalence without analytical depth examination; [33] list barriers without quantifying solution-oriented research. Third, theoretical advancements to transition studies and agricultural development economics are introduced. Whilst [12] and [16] examine technological niches and regime frameworks, they inadequately theorize financial infrastructure as unique transition sector. The ‘structural financial ambiguity’ concept bridges this void, elucidating how epistemic frameworks create and sustain uncertainties they claim to mitigate.

This exceeds beyond [32] descriptive elucidation or [33] barrier listing toward explanatory critique: identification not merely of absences (financial analysis) but of reasons for perpetuation (cognitive oligopoly, methodological entrenchment, North–South epistemic disparities) and implications (climate finance commitments without executable pathways). In institutional economics terms, transaction cost obstacles to CSA finance—information imbalances, enforcement complications, coordination hindrances—remain insufficiently addressed scientifically because the research framework itself embodies these obstacles (English publication costs, Northern journal access fees, CGIAR project timelines). The ‘structural financial uncertainty’ concept introduces theoretical advancements beyond descriptive frameworks, elucidating why plentiful climate finance commitments fail to convert into practical tools: academic discourse itself sustains ambivalence through epistemic frameworks marginalizing financial mechanism investigations.

## 5 Conclusion

The bibliometric analysis enhances transition studies by uncovering financial infrastructure as the critical element in agricultural transformation theory. Whilst Geels’ Multi-Level Perspective focuses on technological niches and regime landscapes, African CSA demonstrates a ‘financial void in the middle’: strong niche innovation (agricultural methods) and ambitious landscape pressures (climate finance commitments) lacking supportive financial mechanisms (lending systems, risk management tools, payment frameworks) necessary for effective translation. The ‘structural financial ambiguity’ identified constitutes not merely literature gap but fundamental aspect of socio-technical systems favoring technical advancements over institutional progress. This conclusion questions underlying beliefs—common in recent assessments—that CSA transition constitutes merely knowledge transfer issue; analysis indicates it is essentially financial infrastructure co-construction problem demanding equivalent epistemic decolonization to technological sharing.

### 5.1 Beyond observation: epistemology of climate finance

The bibliometric investigation uncovers a scientific field in contradictory tension. Climate-smart agriculture across Africa undergoes unprecedented exponential escalation—31.3% annually, 161 publications over eleven years—yet numerical surge conceals profound qualitative stagnation. A fundamental funding deficiency has been pinpointed arising not from disciplinary delays but from constrained epistemic structure: cognitive oligopoly maintained by select pioneering agronomists, regional compartmentalization of scientific communities, and systemic financial mechanism obscuration in semantic frameworks. This entrapment illustrates technical paradigm endurance wherein agricultural transformation is perceived as biological engineering challenge rather than institutional and financial engineering problem.

The contribution extends beyond diagnosis. Through ‘structural financial ambiguity’ introduction, a theoretical framework is provided illustrating how scientific literature, rather than alleviating policymaker concerns, cultivates and perpetuates them. This uncertainty stems not from understanding deficiency—economic obstacles are thoroughly documented—but from incapacity to convert diagnosis into actionable solutions. When 78% of studies highlight financial limitations as primary implementation hurdles yet merely 5.6% link economic evaluations with adoption factors, the academic realm creates unmet expectations: promising change without equipping stakeholders with financing means.

### 5.2 Reconfiguration imperatives

Three mechanisms for field evolution are identified: viewpoint transformation, economics systematization, and political relevance infusion.

#### Viewpoint transformation

Collaboration mapping uncovers uneven scientific landscape skewing research focus. Predominance of activity in Eastern and Southern Africa—where funding frameworks are available, albeit imperfect—distracts from distinct Sahel and Central African requirements, where credit, insurance and environmental service payment system absences present fundamental implementation obstacles. Knowledge redistribution toward these ‘blank areas’ is essential, not from scholarly generosity, but for scientific significance: financial strategies cannot be simply applied across settings without thorough contextual adjustment.

#### Economics systematization

Post-2021 ‘new wave’ emergence—Tabe-Ojong, Antwi-Agyei, Getnet—embodies paradigmatic transformation potential. These scholars employ robust econometric techniques (discrete choice experiments, contingent cost–benefit evaluations, randomized impact assessments) to evaluate financial tool efficacy and acceptance. This economic challenge systematization must become standard practice, not anomaly. It entails discipline blending: agronomy, behavioral economics, climate finance political science, financial innovation sociology. Prominent journals hold editorial obligation to broaden pages to economic analyses without relegating them to trivial concerns.

#### Political relevance infusion

The CSA domain functions as political dimension-stripping device: by framing agricultural change as technical practice adoption challenge, it conceals power dynamics dictating financial resource access. The top 12 authors, primarily associated with international organizations or South African universities, serve not merely as knowledge producers but as gatekeepers influencing research funding direction and, consequently, public policy agendas. Epistemic democratization promotion—empowering emerging African scholars, valuing indigenous informal financing wisdom, incorporating non-academic stakeholders (cooperatives, fintech companies, agricultural banks)— is essential for CSA research to cease perpetuating inequities it purports to address.

### 5.3 Toward financial evolution discipline

Field advancement hinges on capacity to experience financial shift akin to linguistic or ethical transformations reshaping other disciplines. This transformation entails viewing financing not as external factor or contextual limitation, but as pivotal investigation focus: farmer interpretation and evaluation of various financial tools; institutional prerequisites for global climate finance commitment materialization as accessible smallholder capital; hybrid business model (public–private–community) identification guaranteeing CSA practice economic sustainability whilst ensuring disadvantaged inclusion.

These inquiries constitute essential underpinnings, not additional burdens on congested research agendas. Without scientifically validated responses, COP29 commitments, national investment frameworks and adaptation plans remain hollow assurances. Africa does not lack technical advancements; it struggles with epistemically robust financial innovation shortage. The bibliometric assessment, by rendering overlooked elements visible, creates opening. Scientific community expansion into comprehensive paradigmatic shift—or recognition that, in present condition, contribution weighs more toward issues than resolutions—remains incumbent.

Despite regional constraints, the bibliometric exploration uncovers dual injustice: Sahel and Central African areas encounter severest climate threats whilst generating least prominent (English-language, Scopus-indexed) scholarship; concurrently, alternative funding frameworks (solidarity-driven, government-led) remain scientifically unrecorded and omitted from worldwide climate financing structures. Future inquiries must emphasize mixed-language systematic assessments, grey literature incorporation and South–South knowledge collaborations to amend these disparities. The ‘structural financial uncertainty’ identified might, in reality, prove more pronounced in francophone Africa—not from funding absence, but from informal and formal pathway functioning hidden from prevailing bibliometric paradigms.

## Data availability statement

The data supporting this study are derived from the Scopus database, which is a third-party commercial source. The search query and selection criteria are fully described in Section 2.1 and 2.2. The metadata used for analysis (CSV and BibTeX files) are available from the corresponding author upon reasonable request. No original data were generated in this bibliometric study.

## Funding

No funding was received for conducting this study.

## Competing interests

The author has no relevant financial or non-financial interests to disclose.

## Ethics approval and consent to participate

Not applicable

## Consent for publication

Not applicable

## Declaration of AI use

During the preparation of this work, the author used Grammarly, DeepL, Rewrite Guru to improve language readability and assist with paraphrasing complex sentences. After using this tool, the author(s) reviewed and edited the content as needed and take(s) full responsibility for the content of the publication.

## Notes

### Competing Interest Statement

The authors have declared no competing interest.

